# GViNC: an innovative framework for genome graph comparison reveals hidden patterns in the genetic diversity of human populations

**DOI:** 10.1101/2024.06.10.598220

**Authors:** Venkatesh Kamaraj, Ayam Gupta, Karthik Raman, Manikandan Narayanan, Himanshu Sinha

**Affiliations:** Centre for Integrative Biology and Systems Medicine (IBSE), Wadhwani School of Data Science and AI, Indian Institute of Technology (IIT) Madras, Chennai 600036, India; Wadhwani School of Data Science and AI, IIT Madras, Chennai 600036, India; Department of Data Science and AI, IIT Madras, Chennai 600036, India; Department of Computer Science and Engineering, IIT Madras, Chennai 600036, India; Department of Biotechnology, Bhupat and Jyoti Mehta School of Biosciences, IIT Madras, Chennai 600036, India

## Abstract

**Motivation:** Genome graphs represent genetic diversity by highlighting polymorphic regions, but current methods lack the ability to characterize and compare their complex structures effectively.

**Results:** Our study introduces GViNC: a framework for Genome graph Visualisation, Navigation, and Comparison. GViNC maps genomic coordinates onto genome graph nodes, facilitating subgraph partitioning by regions, which aids in navigating and comparing genetic data. Applied to multiple genome graphs from the 1,000 Genomes Project, we observed that genomic complexity varies by ancestry and chromosomes, with rare variants increasing variability significantly. GViNC identified key regions like HLA and DEFB loci, revealing population-specific heterogeneity linked to essential biological functions. Its versatility and scalability support extensive research on genetic diversity across different cohorts or species.

**Availability and Implementation:** GViNC, automated with Snakemake, is available at https://github.com/IBSE-IITM/GViNC.

**Contact:** (K.R.), (M.N), (H.S.)

**Supplementary information:** A supplementary document with tables and figures accompanies this manuscript.

## INTRODUCTION

Genome graphs offer a richer representation of a collection of genomes than the traditional linear reference-based depictions. They are suitable for representing highly heterogeneous populations as they can accommodate multiple genetic variations at a single genomic position [1]. In particular, variational genome graphs encode the genetic variants within a population and embed paths that can represent possible alternate sequences from a population for which the reference is built [2, 3]. They methodically segment sequences into discrete entities termed nodes. The interconnection of these nodes through edges establishes a network, facilitating seamless traversal within the graph from one node to another to capture possible alternate sequences. Genome graphs can incorporate information from multiple individuals from varied populations, making them more suited for studying diverse genomic landscapes [4]. Previous studies broadly classify genome graphs as pan-genome or population-specific. While a pan-genome graph represents the genetic diversity across multiple populations, a population-specific genome graph captures the genetic variations like single-nucleotide polymorphisms (SNPs), insertions, deletions (INDELs), and other variants prevalent within a particular population [5].

The structure of a genome graph mirrors the genetic diversity of the represented samples and makes it possible to conduct comparative studies focusing on hypervariable regions at a population-specific and personalised level. Despite this inherent advantage, methods to comprehensively study the structural complexities underlying genome graphs are limited. In particular, methods to compare the complexity of genome graphs of different populations are lacking. A recent study on quantifying the complexity of a single genome graph used metrics such as the total number of nodes, edges, and connected components as indicators of the complexity of the graph [6]. Although helpful in explaining the overall complexity, these metrics do not represent the intricate and localised complexities intrinsic to genome graphs. So, these metrics cannot be used for a well-grounded quantitative comparison of entire genome graphs. Furthermore, existing genome graph visualisation tools focus on representing fine-scale sequence variations and localised topology, limiting their utility to smaller genome graphs containing a few thousand nodes [7–13]. These tools cannot operate on large graphs with millions of nodes, such as the human genome graphs. There is a need for a comprehensive framework that can bridge the complexity at the level of sequence variation to the topology of whole genome graphs. Such a framework is a prerequisite to navigating and elucidating the genetic diversity of the represented species or population through the lens of genome graphs. The framework can help capture, quantify, visualise, and compare the complexities of genome graphs and shed light on the regions of the genome with salient structural features. It would enable researchers to locate previously unexplored parts of the genome, possibly particular to a population, with potential functional significance.

The study introduces GViNC (Genome graph Visualisation, Navigation and Comparison), a framework that characterises genome graph structures, emphasising sequence variations and genome topology. GViNC visualises entire genome graphs, identifying key loci with significant individual variability and providing metrics for comparing structural complexities. Applied to human genome analysis from the 1,000 Genomes Project, it revealed shared and unique genomic complexity patterns among five populations, influenced by ancestry and chromosomes. The framework highlighted biologically significant regions, such as the HLA and DEFB loci, and identified hypervariable sequences in genes like *MICA* and *NOTCH4*. Overall, we demonstrated our proposed method’s capability, versatility, and scalability to perform in-depth analyses of the genetic complexities within and between human populations.

## METHODS

### Construction of human genome graphs

Recent developments using genome graphs as a reference structure for genome analysis have given rise to numerous tools for their construction [14]. While most of the existing tools work adequately for smaller genomes, they rarely perform optimally as the size of the genome scales up. Very few software programs run smoothly at the scale of human genomes, with *vg toolkit* [2] being one of them. The *vg toolkit* is one of the widely used, openly available tools that can efficiently handle the construction of human genome graphs and analyse whole genome sequences with the same. It is an actively maintained software with an extensive set of functionalities. Based on these characteristics and our aim to use an open-source software framework to develop tools for genome graphs and their analysis, we used the *vg toolkit (version v1.42.0)* in our study to construct human genome graphs.

Genome graph construction with the *vg toolkit* involved incorporating variants over a linear reference structure to generate the edges and nodes that comprised the graph-based reference structure. We have used the GRCH38 (hg38) reference genome as the linear reference structure for constructing the genome graphs. The 1000 Genomes Project [15], built on the Human Genome Project, sequenced hundreds of individuals from diverse populations and revealed millions of genetic variations. The variants from this project (1KGP variants), composed mainly of short variants like SNPs and INDELs, captured the genetic heterogeneity of the species well and were used to construct the human pan-genome and population-specific graphs. As the 1KGP variant set called with the hg38 reference genome was incomplete for chromosome Y, we lifted over the Y chromosome variants from hg37 to hg38 and used them to construct and analyse genome graphs.

The 1KGP variant set was composed of variants called from 2504 individuals from five major ancestries: African (AFR), American (AMR), East Asian (EAS), European (EUR), and South Asian (SAS). The entire 1KGP variant set comprising nearly 85M variants was considered in constructing the human pan-genome graph. The variants called in the samples from each particular ancestry were filtered out and used to construct the respective population-specific genome graph. To discern the effect of rare variants on the complexity of genome graphs, we also created one constructed using only the common variants with an alternate allele frequency of at least 0.05. The details of the input variant set and the names of some of the constructed genome graphs are given in Table 2. It can be noted that the number of variants differs for each population based not only on the sample count but also with respect to the genetic diversity of the samples and their nucleotide divergence from the reference genome. Furthermore, we have also constructed subgraphs specific to those regions for the in-depth study of specific genomic locations from these larger human genome graphs.

### Structural analysis of genome graphs

Genome graphs encase within them a reference path that corresponds to the linear reference genome on which the variants are augmented [2]. The human genome graphs constructed in our study encompassed a path that retraced the hg38 reference genome. Nodes in the reference path backtracked on the linear coordinate system, and this property of the genome graph was a bridge between the two reference structures [14]. Every allele of a variant added to the genome graph created a path that diverged from the reference path at a particular reference node. Each variant path that diverged from a reference node would increase the out-degree of that corresponding reference node by one. This implied that the out-degree of the node directly correlates with the genetic diversity of the input samples at that genomic position. This phenomenon was scaled up to the whole genome and was used to credibly quantify the complexity of the entire human genome through the lens of genome graphs. To systematise the structural analysis of genome graphs, we propose the definitions in Table 1.

**Table 1:**
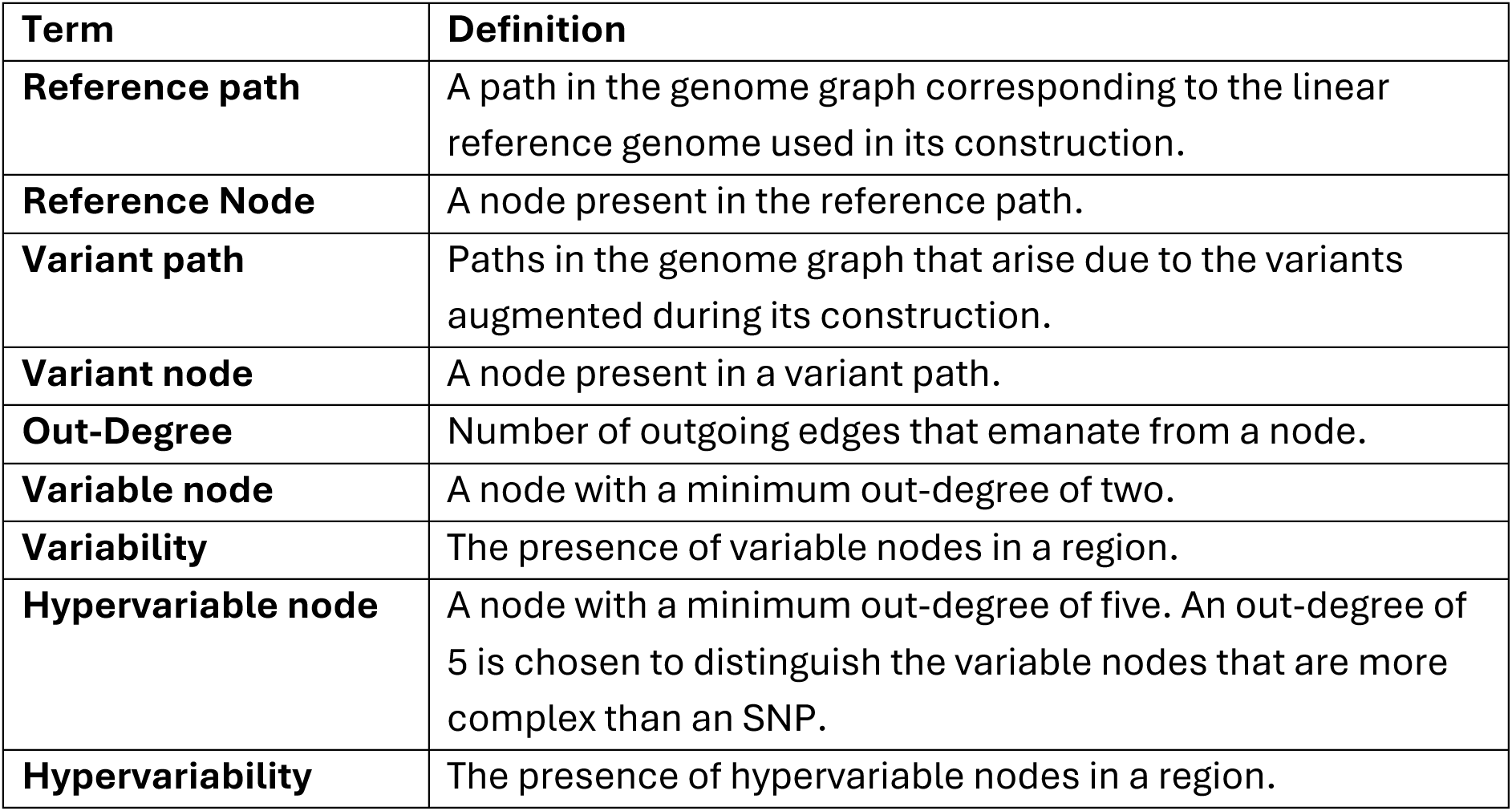
Definitions of genome graph terms used in this paper.

**Table 2:** Description and the number of input variant sets used in the construction of each of the genome graphs constructed in the study. The number of variants and the sample size reflect the genetic heterogeneity of the populations. One individual has dual ancestry of both the African and European population groups.

| No. | Description of Input Variant Set | Number of Samples | Number of Variants | Name of the Genome Graph |
| --- | --- | --- | --- | --- |
| 1 | All the variants from the 1KGP project | 2,504 | 85,123,169 | 1KGP Genome Graph |
| 2 | Variants corresponding to the African ancestry | 662 | 43,757,534 | AFR Genome Graph |
| 3 | Variants corresponding to the American ancestry | 347 | 29,539,124 | AMR Genome Graph |
| 4 | Variants corresponding to the East Asian ancestry | 504 | 24,759,577 | EAS Genome Graph |
| 5 | Variants corresponding to the European ancestry | 503 | 25,196,487 | EUR Genome Graph |
| 6 | Variants corresponding to the South Asian ancestry | 489 | 27,874,233 | SAS Genome Graph |
| 7 | Common variants from the 1KGP variants | 2,504 | 8,496,706 | Common_1KGP Genome Graph |

The genome graphs created in the *vg toolkit* were extracted in the Graphical Fragment Assembly format (GFA) and were incorporated into a custom *Python* program built on top of *NetworkX* [16]. *NetworkX* enabled the effortless application of graph algorithms at the scale of human genomes. We subdivided the reference path in the constructed genome graphs into bins of suitable genomic windows based on the reference genome length. For example, the genome graphs at the scale of the entire human genome were divided into 10 Mbp windows. Out-degrees of all the nodes were calculated for the binned subgraphs from all the genome graphs. Variable and hypervariable nodes from each bin were enumerated. A complete panoramic view of the genome graphs was obtained by collating the count of such nodes throughout the human genome. Circos plots [17] were used to get a bird’s-eye view of the genome graphs. The entire workflow (Figure 1), which consisted of custom scripts in multiple programming languages and other existing open-source tools, was automated using the Snakemake workflow management system [18]. It can be noted that the input data required to run GViNC was only a linear reference genome in FASTA format and a collection of variants in VCF format that was called with respect to the same reference genome. The bird’s eye views of various genome graphs can be used to visually compare them and identify the overall structural similarities and differences.

**Figure 1:**
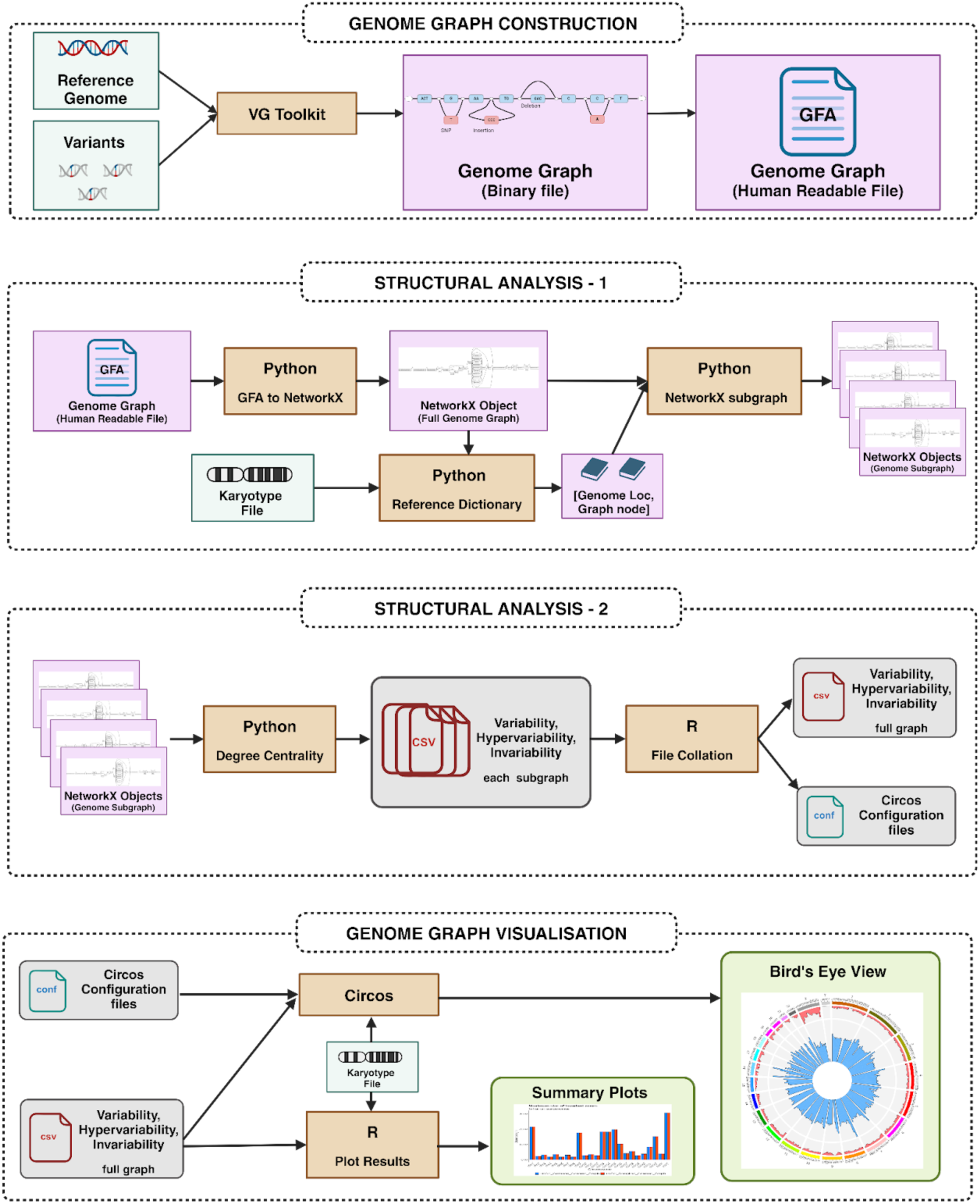
GViNC: A framework for genome graph visualisation, navigation and comparison. Illustration of the steps involved in the GViNC workflow. It can analyse the structural complexities in a genome graph and shed light on the genetic complexity of the samples the graph represents.

The functional significance of crucial findings from the structural analysis of genome graphs was studied in depth. Furthermore, for the quantitative comparison of the structures of genome graphs, we devised the following metrics:

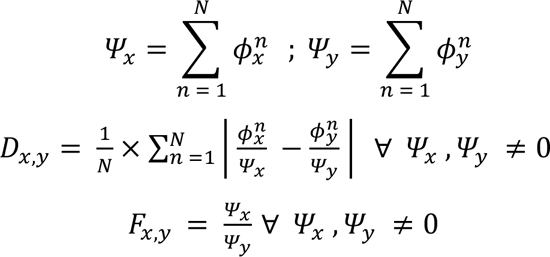

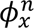: *variability in the genome graph: x in a particular subgraph : n*

𝛹_!_: *The total variability of genome graph x*

*N: Total number of subgraphs. Should be the for both the genome graph*

*D*_!,’_ : *Distance in term of variability between two genome graph x and y*

𝐹_!,’_ : *Fold change of variability between two genome graph x and y*

The distance metric 𝐷_!,#_ takes into account the differences in the input variant sets. This metric effectively found the distance between the two genome graphs, normalising for the total variability, accounting for the difference in input variants size of each genome graph, as well as the total number of subgraphs that were created, accounting for chromosome size differences. As 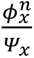 is the probability of a variant occurring in a particular genomic window (or subgraph: n), 𝐷_!,’_ could also be explained as a normalised version of the Total Variational Distance, a widely used measure to find the statistical distance between probability distributions. It can be noted that 𝐷_!,’_ could be leveraged to compare whole genome graphs as well as chromosome-specific and loci-specific genome graphs when it was calculated for only the subgraphs specific to that region. Moreover, the distance metric satisfied the commutative property, making the order of the genome graphs irrelevant. The formulae for distance and fold change metrics illustrated above describe them between two genome graphs in terms of variability. The same formulae were generalised for the distance/fold change between two genome graphs in terms of hypervariability, if the values corresponding to variability were substituted with the values of hypervariability. The formulation of both the distance metrics 𝐷_!,’_ and 𝑭_𝒙,𝒚_ ensures that the resolution of comparison could be adjusted for the genome size. 𝑭_𝒙,𝒚_, being the fold-change would be useful for comparing the larger shifts in the structures for a pair of genome graphs, ideally constructed with the same number of samples. 𝐷_!,’_ could even identify the subtle changes in heterogeneity between two cohorts of samples, and the normalisation factors in its formula account for differences in the cohorts’ sample sizes.

## RESULTS

### GViNC: The framework for the structural analysis of genome graphs reveals genetic diversity

Genome graphs capture the complexity and diversity of a species, and analysing its structure can be crucial for understanding specific aspects of its genome. However, the structure of a genome graph is dynamic and depends on the set of variants augmented during its construction. GViNC is designed to effectively quantify the genomic complexity encompassed by the genome graph from the perspective of variability and hypervariability. Each allele of the variant added to the genome graph formed a distinct path that branched off from the reference path at a specific reference node, increasing the degree of that particular node by one. This meant nodes with an out-degree of at least two originated at the sites of variants in the genome. So, the number of variable nodes in a genome graph characterised the number of variants augmented during its construction.

The maximum possible out-degree of a node representing an SNP in the genome graphs was four (one reference and three possible alternate alleles). Intending to capture the highly complex parts of the genome, we defined hypervariable nodes as more complex than SNP nodes. Hypervariable nodes, representing genomic regions with high levels of polymorphism, could be essential for studying population-specific genetic diversity and capturing regions of the genome that substantially impact various biological processes and functional relationships. These hypervariable nodes could be used as markers to distinguish different genome graphs built for the same species, as other graph-derived metrics like the number of nodes and edges cannot achieve this.

In our study, we constructed multiple pan-genomic and population-specific human genome graphs and performed structural analysis on each. We enumerated the number of variable nodes and hypervariable nodes in the defined subgraphs. Variability and hypervariability of the genome graph were quantified by the total counts of variable and hypervariable nodes in the subgraphs, respectively. This opened up the opportunity for detailed visualisation of the genetic diversity, at the scale of the human genome, for a large number of samples in a single figure.

### Visualisation of 2,504 human genomes in a single figure

The human pan-genome constructed with the 1KGP variants captured the genetic diversity of 2,504 individuals. The reference path in this genome graph, corresponding to the hg38 reference genome, was subdivided into 10 Mbp tiled bins. A subgraph was extracted from the pan-genomic graph for each defined bin. Then, the prevalence of variable and hypervariable regions in each bin in each chromosome was obtained. The variability and hypervariability observed in every subgraph were used to get a bird’s-eye view of the entire genome graph, capturing the genomic complexity of 2,504 humans in a single figure, illustrating a panoramic view of the human pan-genome graph (Figure 2).

**Figure 2:**
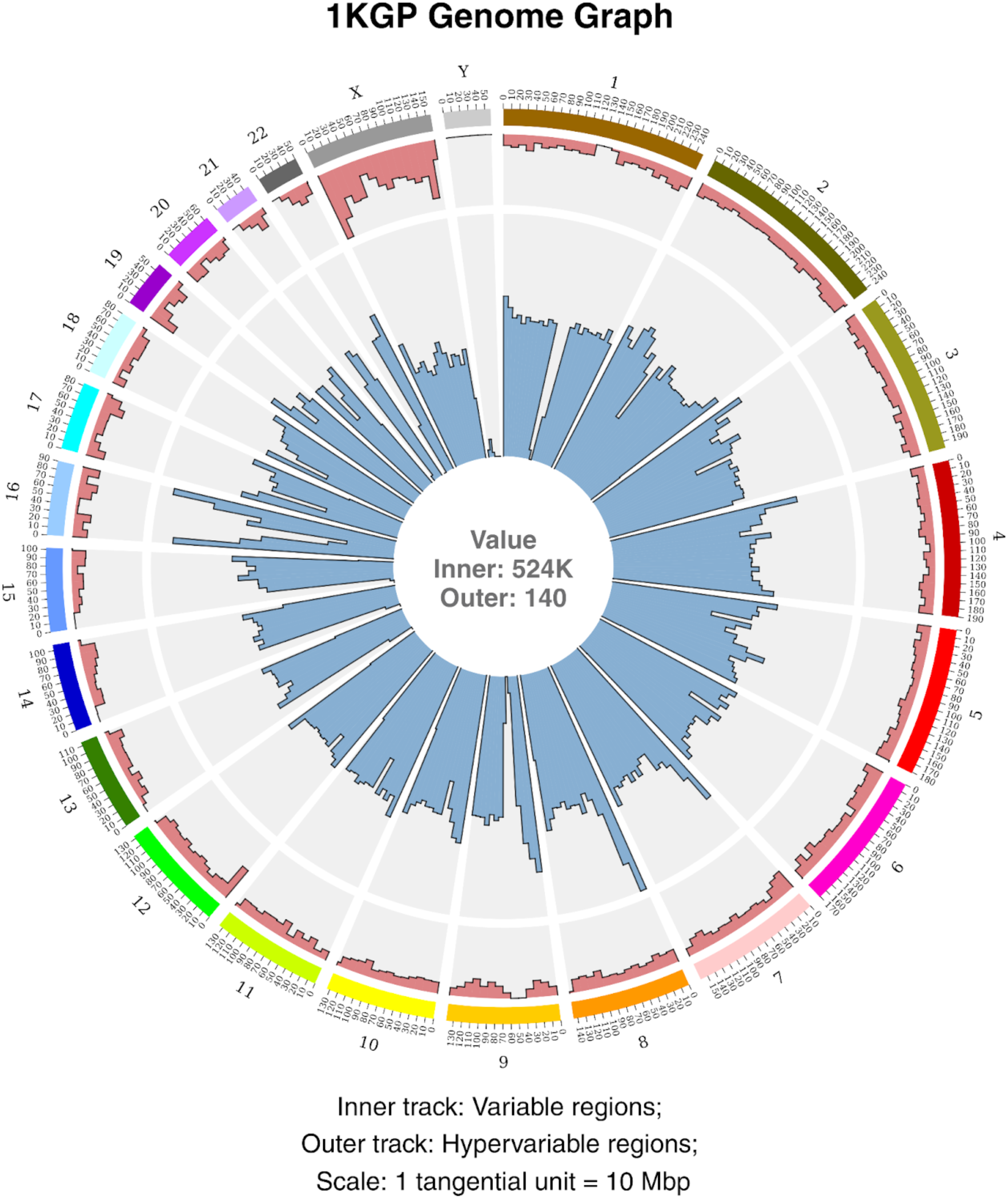
Bird’s-eye view of human pan-genome graph. A single-figure representation of an entire genome graph, capturing the genetic diversity of thousands of human samples spread across multiple populations. There are 24 sectors: one for each of the 22 autosomes, one for chrX, and one for chrY. The subgraph size (tangential scale) for the genome graphs was set as 10 Mbp. The number of hypervariable nodes in each bin or the extracted subgraph is depicted in the outer track. The number of variable nodes in each bin is represented in the inner tracks. The text at the centre indicates the maximum value for the inner and outer tracks.

The highest variability of 524K was observed in chr8 for the human pan-genome graph. High variability was also observed in certain regions in chr16. A closer look at these high variability zones will be taken up in the latter part of this study. Even though a majority of the hypervariability was present in chrX, hypervariable nodes were observed throughout the genome, albeit in smaller numbers. Out of the total 7,446 hypervariable nodes observed, two nodes were seen with the maximum out-degree spanning twelve. When the exact location of the most hypervariable nodes was determined, it was found that both these nodes were adjacent to each other and were present in chr1. Figure S1 depicts the path-level representation of the adjoining 12-degree nodes in the 1KGP Complete genome graph obtained using the default visualisation command bundled with the *vg toolkit*.

GViNC, apart from achieving single-figure visualisations of large numbers of samples, irrespective of the scale of the genome, had additional useful features. The prospect of comparing different genome graphs arose from visualising entire genome graphs and quantifying variability and hypervariability in them. A visual inspection could be done to compare and uncover genomic regions that were unique or different for sample sets or populations. On the other hand, quantitative comparison using variability/hypervariability could aid in identifying populations that might be close to or farther from each other in terms of genetic diversity. We demonstrated such comparisons using the population-specific genome graphs constructed from defined subsets of the 1KGP samples.

### Population-specific genome graphs unveiled shared and unique genetic signatures for different human populations

The panoramic visualisations of five population-specific genome graphs were obtained using GViNC (Figure 3A-E). Even though the bird’s eye views were scaled to the same size to make visual comparisons easier, it could be inferred from the maximum values at the plot centre that the AFR genome graph represented more individuals. The maximum values follow the trend of the total number of input variants (Table 2). We saw an increased variability in all populations for certain bins from chromosomes 4, 8, and 16, similar to the observation in the human pan-genome graph (Figure 2). We could also observe that the hypervariability was concentrated in chrX in all five populations, as observed in the case of the pan-genome. However, the number of hypervariable nodes in chrX in individual populations was much less than that of the pan-genome, as reflected in the maximum values. While the first 10 Mbp bin in the human pan-genome contained 140 hypervariable nodes, the same bin only contained 97 such nodes in the AFR genome graph and dropped to 40 in the EUR genome graph.

**Figure 3:**
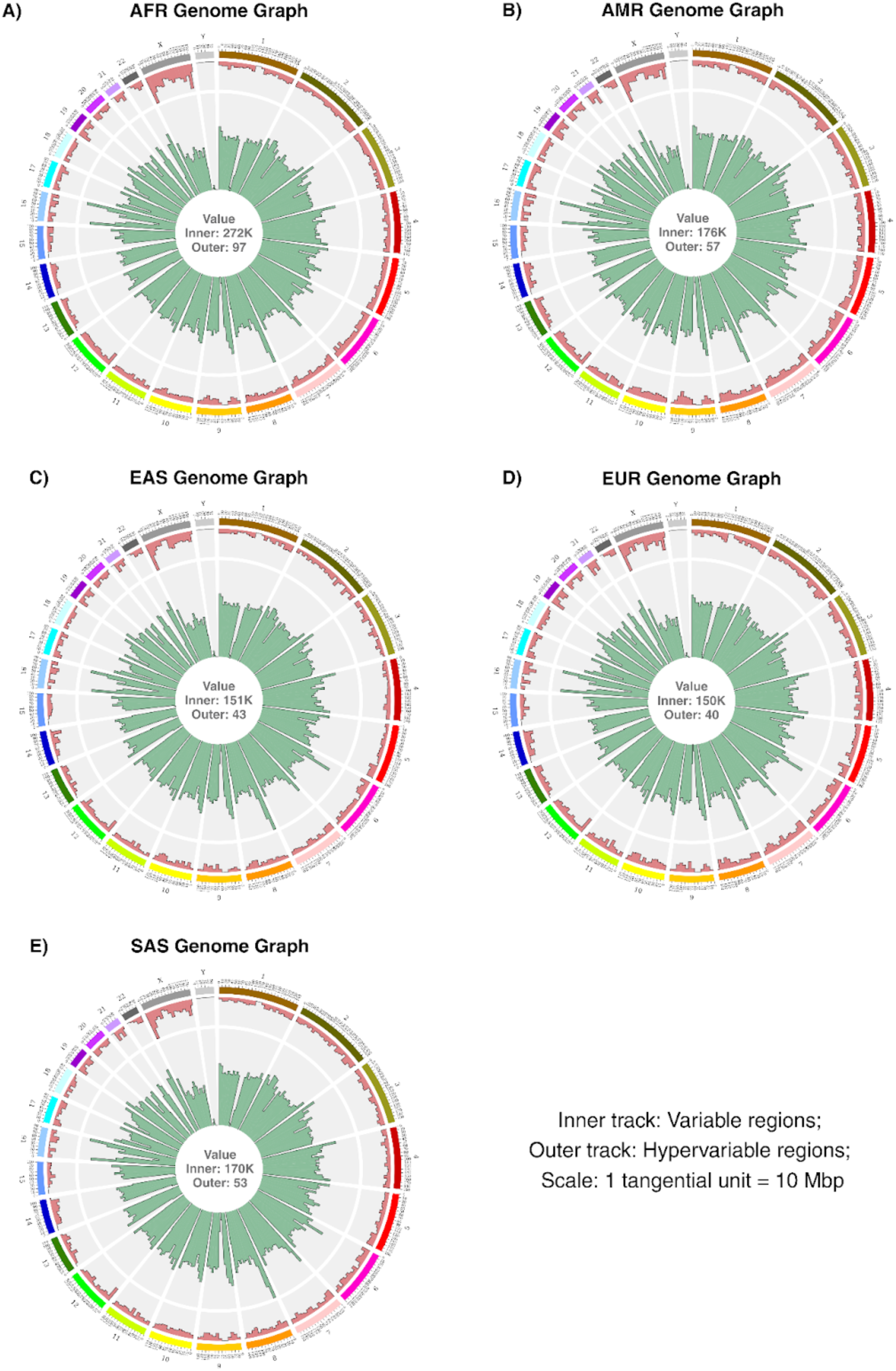
Bird’s eye view of population-specific human genome graphs. Single-figure representations of population-specific genome graphs. The text at the centre of each plot indicates the respective maximum value for the inner and outer tracks. The tangential scale (subgraph size) for all the genome graphs was 10 Mbp. **(A)** African Genome Graph, **(B)** American Genome Graph, **(C)** East Asian Genome Graph, **(D)** European Genome Graph, **(E)** South Asian Genome Graph.

Upon careful inspection, we could identify subtle differences in the contours of the bars in variability and hypervariability between the five populations. However, visual comparison of more than two genome graphs at a time was a laborious and subjective process. To overcome this problem and to get a quantitative readout for the comparisons of genome graphs, we calculated the graph distance 𝐷_!,#_ between all ten pairwise combinations of the five populations, represented as heatmaps [19] of these pairwise distances in terms of both variability (Figure 4A) and hypervariability (Figure 4B). We have also identified the top three chromosome clusters using Euclidean distance-based hierarchical clustering.

**Figure 4:**
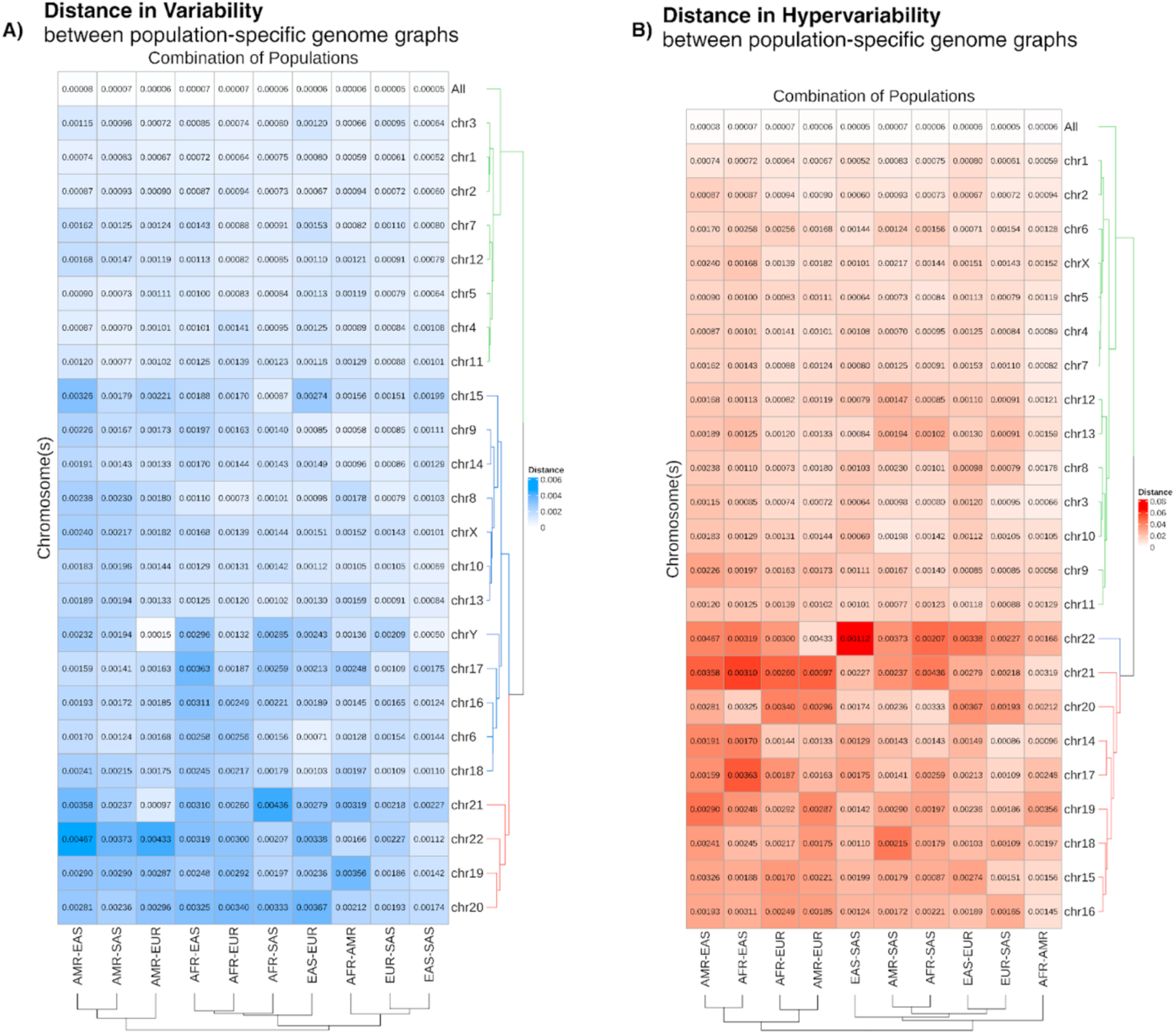
Distance between population-specific human genome graphs. **(A)** 𝐷_!,’_calculated for variability in all possible pairwise population combinations. The metric was calculated for the whole genome (shown as All) and individual chromosomes. **(B)** 𝐷_!,’_ calculated for hypervariability in all possible pairwise population combinations. As hypervariability is completely absent in chromosome Y, we cannot calculate the distances as 𝐷_!,’_ would be mathematically indeterminate.

The distance between populations was minimal for both variability and hypervariability when calculated for the entire genome. The distances were higher when calculated over individual chromosomes. The cluster with high distance metrics across population combinations contained chromosomes 19, 20, 21, and 22 for variability and only 22 for hypervariability. It could be noted that these were some of the smallest autosomes present in the human genome, and they generally showed higher 𝐷_!,#_, indicating these chromosomes had more genetic differences between populations compared to the whole genome. The larger autosomes clustered together with the whole genome in both variability and hypervariability, showing reduced distance between different population combinations. ChrX clustered together with the larger autosomes for hypervariability distances, but not in variability distances. ChrY, being a short chromosome, did not cluster with the smaller autosomes in variability distance. Taken together, the quantitative comparison of population-specific genome graphs showed that the genetic diversity of the human populations is not only dependent on ancestry but also varies across chromosomes and seems to depend on length, recombination rates, and other properties of the chromosomes (see Discussion).

### Rare variants sustain variability but drive hypervariability

GViNC allowed for the visualisation and comparison of genome graphs. It could also help in understanding the genetic heterogeneity of different populations. GViNC could also be leveraged to comprehend the effects of rare variants on the genetic diversity of the species or populations. To illustrate this, we created and analysed a human pan-genome with only the common variants from the 1KGP samples and compared it with the complete human pan-genome containing both common and rare variants. A bird’s-eye view was obtained for both the 1KGP Genome Graph (Figure 5A) and the Common_1KGP Genome Graph (Figure 5 B), highlighting the similarities and differences in their variability and hypervariability.

**Figure 5:**
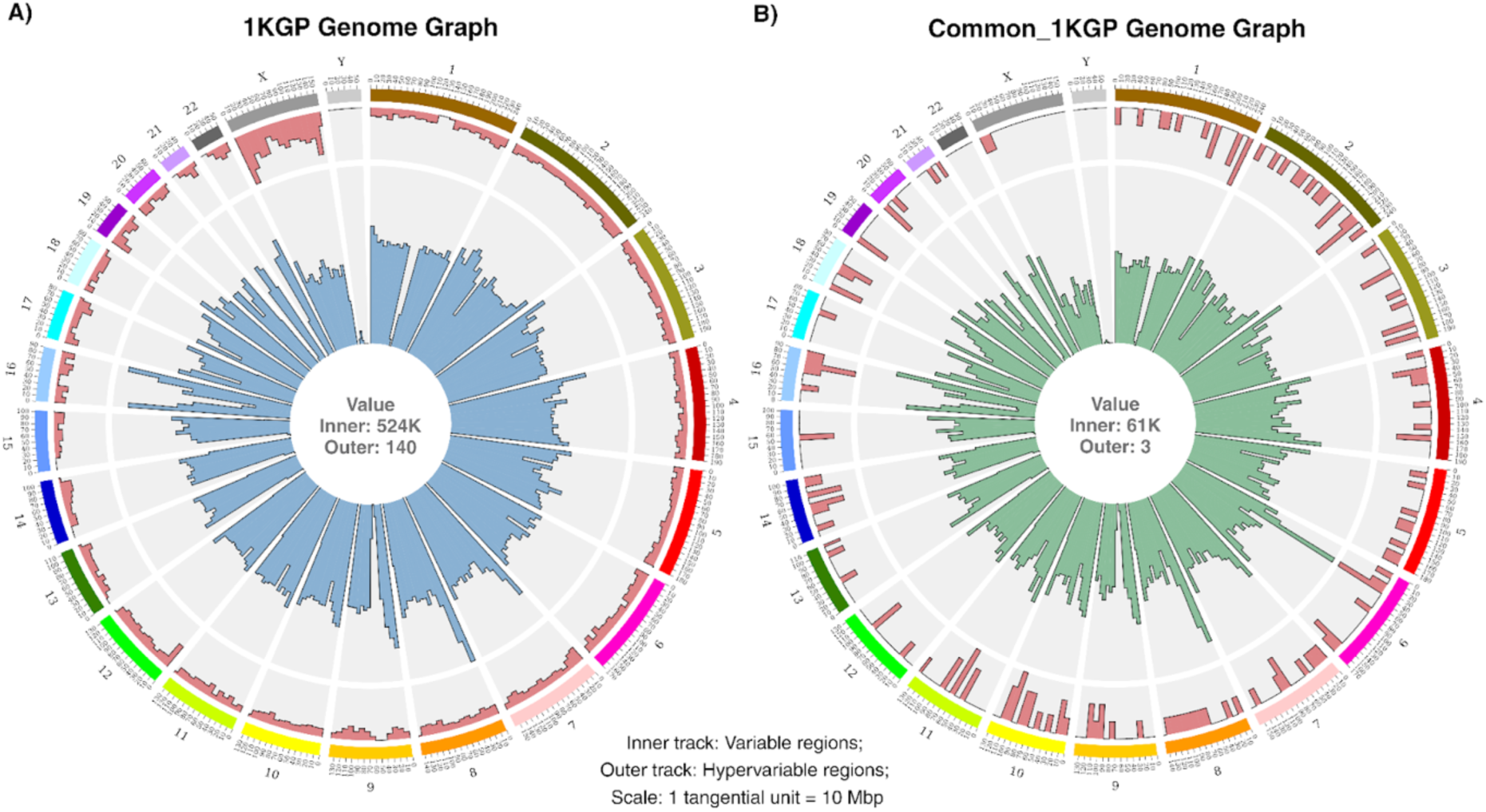
Visual comparison of human pan-genome graphs. **(A)** Single figure representation of the 1KGP Genome Graph that augmented all variants. **(B)** Single figure representation of the Common_1KGP Genome Graph that augmented the variants with an alternate allele frequency of at least 5%. The text at the centre of each plot indicates the maximum value for the inner and outer tracks. The tangential scale (subgraph size) for both the genome graphs was set as 10 Mbp.

Interesting structural properties from the human pan-genome graphs could be visually inferred from this comparison. The inner track, representing variability, was capped at 524K for the 1KGP Genome Graph, whereas it was 61K for the Common_1KGP Genome Graph. This result was expected as the rare variants were removed in constructing the Common_1KGP Genome Graph, reducing the number of variable nodes. However, the contours of the bars in the inner track were similar for both. The Pearson correlation coefficient for the count of variable nodes in the corresponding bins of both the genome graphs was 0.94, indicating a robust linear relationship in their genomic variability. This further implied that the polymorphism in the human genome that arose due to common variants alone retained certain genome topography from the polymorphism that emerged from the entire set of genetic variants.

However, the distribution of the bars in the outer track capturing hypervariability was starkly different for both the genome graphs. A Pearson correlation coefficient of 0.11 and a Spearman correlation coefficient of 0.21 were observed for the count of hypervariable nodes in the corresponding bins of both the genome graphs, indicating a poor relationship in their hypervariability patterns. In the 1KGP Genome Graph, hypervariable nodes were present throughout, with the majority of them located in chrX. Meanwhile, the Common_1KGP Genome Graph lacked hypervariable nodes all over the genome, even in chrX. Therefore, it seemed that rare variants, which were removed from the Common_1KGP Genome Graph, were mostly driving the hypervariability in the genome. This could mean that the hypervariability observed in the human genome, particularly in chrX, originated from different populations independently, as the common genetic variants present across populations did not cause the structural complexity to the same level.

The effect of rare variants on the structural complexity was computed as the fold change 𝑭_𝒙,𝒚_ in the corresponding complexity types (Table 3). It can be noted that the variability of the human pan-genomes represented the count of variants used in their construction, and it closely followed the variant set sizes observed for 2,504 samples (Table 2). The count of variable nodes was higher than the count of variants due to the possibility that alleles of INDELs with similar sequences were simplified, giving rise to more variable nodes. The 10-fold change in variability caused by the rare variants also aligned with the fact that nearly 90% of the variants observed in the 2,504 samples were rare. It was also observed that rare variants had a greater effect on hypervariability, resulting in a 50-fold change between the two genome graphs.

**Table 3:**
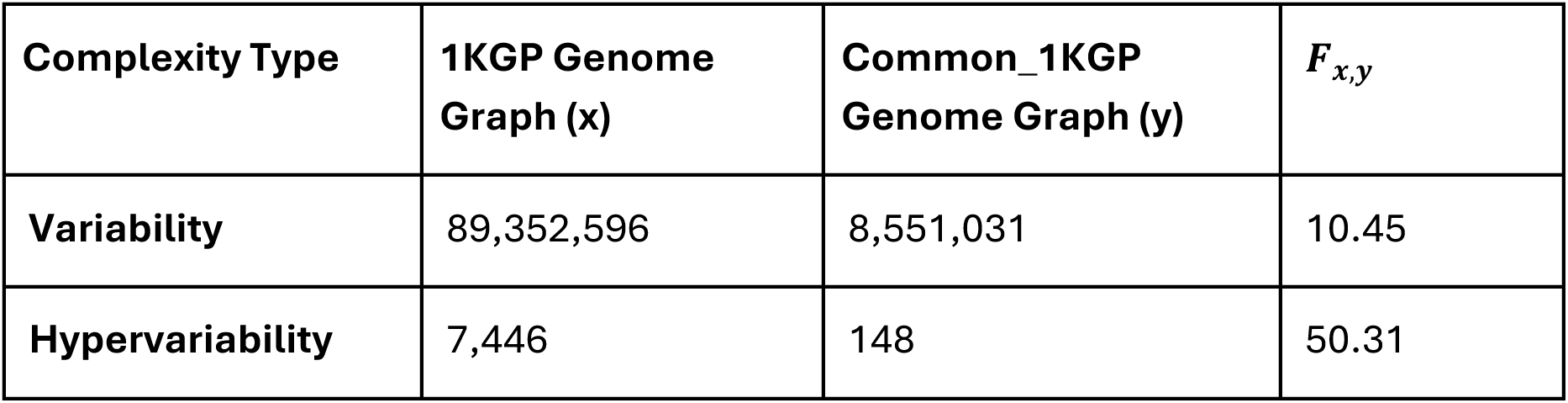
Measures of variability and hypervariability in both the human pan-genome graphs. 𝑭_𝒙,𝒚_is the fold change observed for variability/hypervariability between the genome graphs x and y

### Structural analysis of genome graphs highlights biologically significant loci

GViNC, apart from capturing the genetic diversity of a cohort of samples through genome graphs, also provided a platform for systematic exploration of genomic sites that exhibit high heterogeneity. Such an overview could help us navigate the whole genome of a species to uncover locations in the genome that are highly heterogeneous. When navigation was done on genomic regions associated with fundamental biological functions, we could shed more light on their heterogeneity and prioritise specific locations within these regions that might need in-depth research in the future. While navigating the structure of pan-genomes could reveal high-heterogeneity sites that were common for the species, navigating the structures of population-specific genome graphs could accentuate unique genetic signatures.

We observed that some genomic bins with a high frequency of variable nodes in the two human pan-genome graphs were associated with fundamental functions (Table S1). A 10 Mbp bin in chr6 from 30-40 Mbp showed the highest variability only in the Common_1KGP Genome Graph. This bin encompasses most of the Major Histocompatibility Complex (MHC) region, also known as the Human Leukocyte Antigen (HLA) region. Previous studies have established that the HLA region contains one of the most polymorphic gene clusters of the entire human genome [20, 21]. The presence of multiple HLA alleles in the population ensures that at least some individuals within a population will be able to recognise protein antigens produced by any microbe, thus reducing the likelihood that a single pathogen can evade host defences in all individuals in a given species [22]. However, compared with other bins, the bin containing HLA had a relatively lower frequency of variable nodes in the 1KGP Genome Graph than in the Common_1KGP Genome Graph. This implied that a notable portion of the variants driving polymorphism in HLA were not rare and shared by individuals in different human subpopulations, indicating that many of these variants were ancient [23–25].

In the human pan-genome graphs, other genomic regions with high variability were observed in chromosomes 8 and 16. The first 10 Mbp bin from chr8 with the highest variability in the 1KGP Genome Graph encompassed parts of the β-defensin gene clusters called DEFB. DEFB was previously known to be highly variable and was associated with evolutionary importance and disease risk [26]. Chr16 enclosed a couple of highly variable bins in both human pan-genome graphs as it hosted several large polymorphisms, often associated with segmental duplications [27]. As these bins were highly variable in both the genome graphs, it implied that the polymorphism in these regions originated from common and rare variants. While some of the genomic regions highlighted by this diversity analysis coincided with previously known zones of biological significance, the remaining regions were listed as novel candidate loci whose diversity could be analysed further to ascertain their importance (Table S1, Figure 5).

### Genome graphs of specific loci reveal population-specific heterogeneity in sites of biological significance

The population-specific behaviour of highly polymorphic regions can be deciphered using GViNC. To demonstrate this ability, we constructed population-specific HLA genome graphs at higher resolution than the genome-wide ones and used GViNC to navigate the region and identify interesting zones. The HLA region ranges from the genomic coordinates 29,602,228 to 33,410,226 on chr6 in the hg38 version of the human genome. The input variants used to create each of the HLA genome graphs constructed for the five populations are described in Table S2. The bird’s eye view of the graphs (Figure 6A-E) and the path-level visualisation of some biologically significant loci (Figure 6F-H) were generated. The maximum variability found in each of the graphs was similar for all the populations, even though more input variants were added to the AFR HLA genome graph. The HLA region, one of the highly polymorphic regions in the genome, was not densely populated with hypervariable nodes. However, population-specific patterns were found in the distribution of hypervariability.

**Figure 6:**
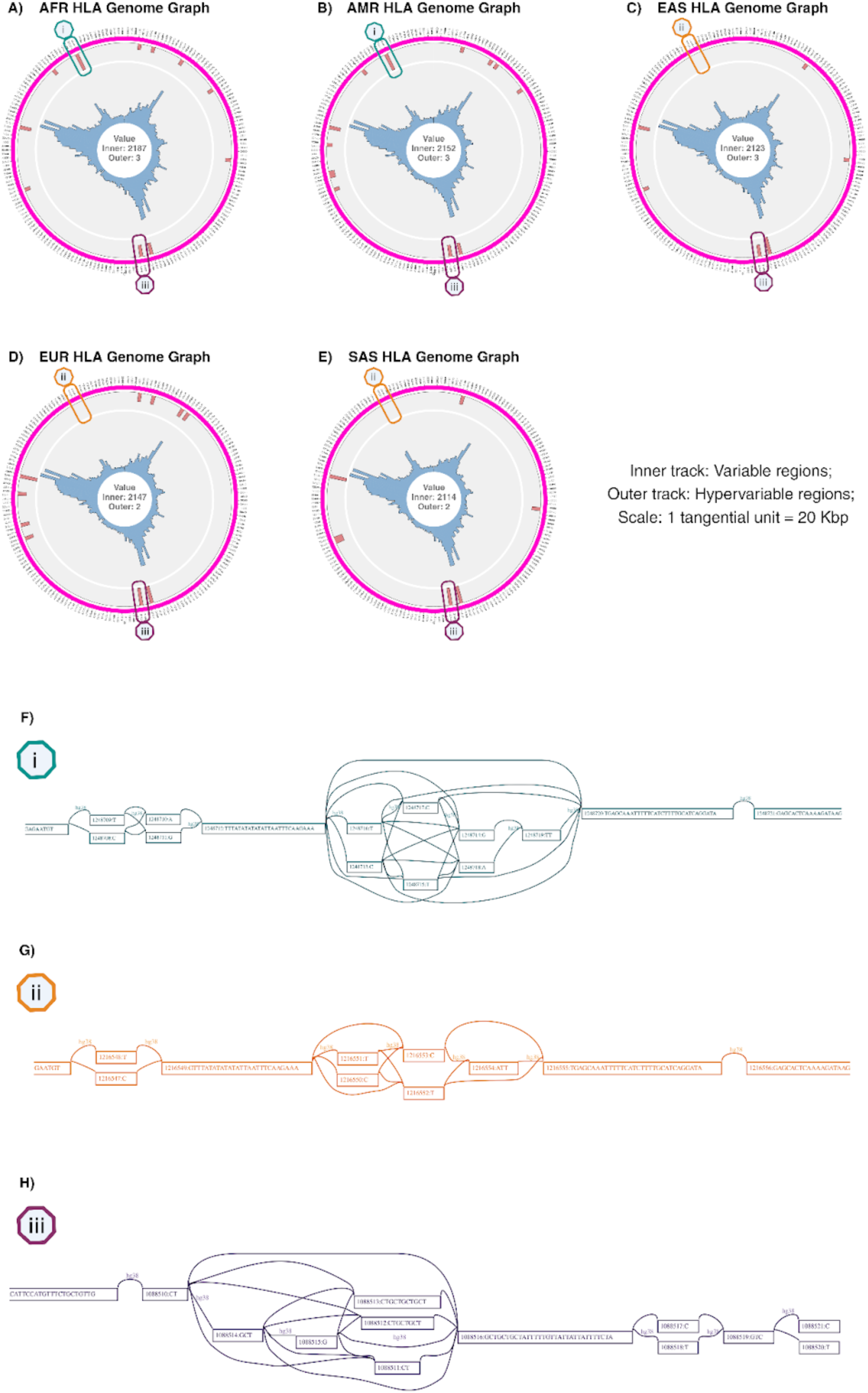
Bird’s eye view of population-specific HLA genome graphs, with the path level representations of *MICA* and *ENSG00000291111*. (A-E) Single-figure representations of HLA genome graphs constructed for African (AFR), American (AMR), East Asian (EAS), European (EUR), and South Asian (SAS) populations. The text at the centre of each plot indicates the respective maximum value for the inner and outer tracks. The tangential scale (subgraph size) for all the genome graphs was 20 Kbp. Markers named (i), (ii), and (iii) are placed on the bins encompassing the *MICA* gene and the *ENSG00000291111* lncRNA. **(F)** Path-level visualisation of the AFR and AMR genome graphs of the HLA region centred around the genomic position 33,127,190. **(G)** Path-level visualisation of the EAS, EUR, and SAS genome graphs of the HLA region centred around the genomic position 33,127,190. **(H)** Path-level visualisation of all the HLA genome graphs centred around the genomic position 31,412,380.

The exact genomic locations of all 19 hypervariable HLA nodes and their occurrence in the population-specific HLA genome graphs are listed in Table S3. To ascertain the functional impact of these hypervariable nodes, we matched these positions with the GENCODE 47 [28] database. We found that, except for one, all the hypervariable HLA nodes were present in genes (Table 4). Seven of these nodes were present in protein-coding genes, and the remaining eleven were found in long non-coding RNAs. Three hypervariable locations were in the protein-coding sequences from exons of the *MICA* and *NOTCH4* genes. One hypervariable node was found in an intron of the *PPP1R10*/*PNUTS* gene, and one in the nonsense-mediated decay region of the *HLA-DRB1* gene. As these regions were functionally significant [20, 29], being hypervariable indicated that individuals within a population have different alleles, increasing the heterogeneity of the particular region’s function.

**Table 4:** Hypervariable nodes in population-specific HLA genome graphs. The genomic position, the type, and the name of the gene are described with additional remarks on the gene’s significance and the population-specific presence of hypervariability.

| Hypervariable Genomic Position | Gene Type | Gene name | Remarks |
| --- | --- | --- | --- |
| 29,699,787 | lncRNA | *ENSG00000310484* | Present only in AFR and EUR |
| 29,767,771 | lncRNA | *HLA-F-AS1* | Promotes colorectal cancer metastasis [44]<br>Present in all except EAS |
| 29,951,575 | lncRNA | *POLR1HASP* |  |
| 29,991,587 | lncRNA | *POLR1HASP* |  |
| 30,182,970 | - | - | Present only in AFR and AMR |
| 30,605,860 | Protein coding | *PPP1R10* | Part of a **retained intron**<br>Involved in DNA repair and apoptosis [45] |
| 31,340,514 | lncRNA | *ENSG00000298396* |  |
| 31,340,587 | lncRNA | *ENSG00000298396* | Present only in EAS |
| 31,412,380 | Protein coding | *MICA* | Both positions are part of a **coding sequence in an exon**<br>A critical component of the immune system.<br>Associated with cancer and autoimmune disorders. [46] |
| 31,412,383 | Protein coding | *MICA* |  |
| 32,211,594 | Protein coding | *NOTCH4* | Present only in SAS |
| 32,223,881 | Protein coding | *NOTCH4* | Is part of the **coding sequence in an exon** |
|  |  |  | Involved in immune regulation and angiogenesis [47] |
| 32,303,314 | lncRNA | *TSBP1-AS1* | Present in AMR and EUR |
| 32,332,596 | lncRNA | *TSBP1-AS1* | Highly expressed in immune cells [48]<br>Present only in AMR |
| 32,492,191 | lncRNA | *ENSG00000307923* | Present only in AMR and EUR |
| 32,589,513 | Protein coding | *HLA-DRB1* | Involved in **nonsense-mediated decay**<br>Plays a crucial role in the immune system and is strongly associated with autoimmune disorders [49] |
| 32,964,843 | Protein coding | *HLA-DMA* | Is part of an **undefined coding sequence**<br>Plays a critical role in antigen presentation and immune regulation [50]<br>Present only in AFR and AMR |
| 33,127,190 | lncRNA | *ENSG00000291111* | Present only in AFR and AMR |
| 33,127,191 | lncRNA | *ENSG00000291111* |  |

Moreover, some of these hypervariable nodes showed population-specific presence. For example, the hypervariable node from the *HLA-F-AS1* non-coding RNA was present in all populations except EAS, and the non-exonic hypervariable position in *NOTCH4* was present only in SAS. The bird’s eye view representations of the five HLA genome graphs showed distinct distributions for hypervariability (Figure 6). When we focused on the *ENSG00000291111* lncRNA at the genomic position 33,127,190, we found that it is hypervariable in the AFR and AMR populations (Figure 6F) while it was not in the others (Figure 6G). However, when we focused on the *MICA* gene at position 31,412,380, we found that all the genome graphs were hypervariable (Figure 6H). Such population specificity implied that the patterns of heterogeneity for these functionally significant zones were sometimes unique to certain populations and sometimes could be shared among different populations.

Finally, we note that this analysis also demonstrated that GViNC is a scalable framework that could not only handle large sample sizes but also analyse single as well as multiple chromosomes – for instance, the bird’s-eye view contained only one section for chr6 in the HLA genome graphs. In contrast, there were 24 sections in the pan-genome graph, one for each chromosome.

## DISCUSSION

GViNC is versatile and allows for an in-depth study of a collection of genomes irrespective of the sample size. It is highly scalable and can even be used to effectively analyse genome graphs that constitute the entire human genome. GViNC can quantitatively compare multiple genome graphs and unearth sites of increased individual-to-individual heterogeneity that can be potentially functionally significant. With increasingly large-scale sequencing studies [30–34], we believe that GViNC strengthens the expanding suite of genome-graph-based tools [2, 7–14, 35–38] and can help the research community perform in-depth examinations and derive fundamental biological insights.

GViNC contains a scalable visualisation technique that can capture the genetic diversity of thousands of individuals and even represent the entire human genome graph in a single figure. Such a panoramic view could act as a signature for a genome graph, capturing the genomic complexity of the underlying set of samples. The single-figure representations of genome graphs also provide the foundation for navigating and prioritising sites of interest, proving valuable for in-depth research.

We devised original quantitative approaches to compare the genetic diversity of different cohorts of samples through the lens of genome graphs. We formulated two metrics that can quantitatively compare the structural complexities of any two genome graphs. In our study, we have used the fold change 𝑭_𝒙,𝒚_to uncover the effect of rare variants on genomic complexities. We found that rare variants increase variability by 10-fold, as expected, and drive hypervariability by a larger 50-fold, indicating rare variants sustain variability and drive hypervariability. We used a custom metric 𝐷_!,’_ to compare the genetic diversity of five human populations at the whole genome level as well as the individual chromosome level. We can observe a trend where the distance 𝐷_!,’_ between genome graph pairs decreased with increasing chromosome length, indicating the dependence of genomic complexity on the chromosome length (Figure 4). The general decrease in complexity with an increase in genome size could be because smaller chromosomes have higher recombination rates than larger chromosomes [39, 40]. These differences set different evolutionary trajectories for chromosomes based on their size, contributing to the difference in genetic diversity [41]. Moreover, the median recombination rates (Figure S2) for human chromosomes generally increase with decreasing length [40]. These observations support the genome diversity-length dependence hypothesis, which was formulated based on our quantitative comparisons of genome graph structures. We strongly believe that GViNC has the potential to significantly impact the research community by catalysing hypothesis-driven analysis, as demonstrated in the examples above.

Apart from providing a platform for systematic comparisons of genome graphs, GViNC also helps researchers navigate them and identify regions with distinct genetic complexities. These regions could potentially be from functionally significant loci. Known regions in the human pan-genome graphs, like the HLA and DEFB gene clusters, and a few novel regions that showed high individual-to-individual variability, were identified by GViNC, demonstrating its ability. Moreover, GViNC also identified functional and novel hypervariable regions in human genomes. Hypervariable positions are those zones in the genome that have greater diversity within the individuals in a cohort. Identifying such regions in a population-specific manner in genomic loci associated with biologically fundamental functions indicates that divergent and variable evolutionary processes in these regions could have resulted in higher divergence between individuals. We also observed that the patterns of hypervariability in these fundamental regions are sometimes shared and sometimes unique to certain ancestries, indicating the population-specific evolutionary trajectories. Furthermore, polymorphic regions in the genome, which vary between populations, can influence disease susceptibility and drug response in different ways. Identifying these regions is key to advancing personalised medicine [42]. Thus, GViNC can benefit both fundamental biological research and the development of strategies for identifying genetic disease loci.

GViNC is coded for and tested on the genome graphs constructed with the *vg toolkit*. However, the idea can be extended to variational genome graphs built with other software, as the structural analysis framework is pivoted only around the traversal of the reference path. GViNC requires minimum inputs to function effectively: a reference genome and a set of variants. In this study, we have used hg38 instead of the latest T2T [43] as the reference genome on which the variant paths were added to create the genome graphs. GViNC would function well, irrespective of the reference genome. Moreover, this proposed idea is not just limited to human genome graphs but can extend smoothly to genome graphs representing the genetic complexity in other species. We showcased the benefits of GViNC using human genome graphs from the 1000 Genomes variants. By applying similar analyses to understudied populations, we can capture genomic complexity, quantitatively compare genetic diversity, and potentially identify key genomic regions that influence diversity, health, and evolution.

GViNC characterises genome graph complexity based on node degree centrality. The annotation of genome graphs with meta-information can complement the idea behind the framework well and would enable the application of more sophisticated graph algorithms to genome graphs. Employing these methods to more diverse genome graphs allow for a comprehensive assessment of genetic diversity, population structure, disease associations, and evolutionary dynamics within and across populations. This information can enhance our understanding of genetic diversity and how it impacts health, diseases, and the historical dynamics of populations.

## DECLARATIONS

## Authors’ contributions

V.K., K.R., M.N., and H.S. conceptualised the study; V.K. developed the computational pipeline and the software; V.K. and A.G. analysed the data; V.K. wrote the first draft of the manuscript; V.K., K.R., M.N., and H.S. critically revised and edited subsequent manuscript drafts; all authors had full access to the data and reviewed and approved the manuscript.

## Acknowledgements

We thank Harshita Agarwal, Veerendra Gadekar, Centre for Integrative Biology and Systems Medicine (IBSE), and Wadhwani School of Data Science and AI (WSAI) members for their valuable discussions and comments.

