## Supplementary Info for "GViNC: an innovative framework for genome graph comparison reveals hidden patterns in the genetic diversity of human populations"

**SUPPLEMENTARY TABLES**

**Table S1:** Top ten bins with the most variability in the human pan-genome graphs. The 1KGP Genome Graph is abbreviated by ‘Complete’, and the Common_1KGP Genome Graph is abbreviated as ‘Common’. Variability is calculated as the count of variable nodes in the corresponding bin.

| Genome Graph | Rank | Region | Variability | Remark |
|---|---|---|---|---|
| Complete | 1 | chr8:0-10 Mbp | 523,905 | Contains parts of the β-defensin gene clusters |
| Complete | 2 | chr16: 80-910 Mbp | 492,481 | Prone to structural variations and duplications |
| Complete | 3 | chr16:0-10 Mbp | 477,967 | Prone to structural variations and duplications |
| Complete | 4 | chr8:10-210 Mbp | 457,413 | Associated with an immune response (BLK) |
| Complete | 5 | chr7:0:110 Mbp | 429,860 | - |
| Complete | 6 | chr9:0:110 Mbp | 426,628 | - |
| Complete | 7 | chr4:0:110 Mbp | 412,151 | Associated with neurological development (HTT) |
| Complete | 8 | chr16:70:810 Mbp | 400,705 | Prone to structural variations and duplications |
| Complete | 9 | chr9:10-210 Mbp | 396,851 | - |
| Complete | 10 | chr3:0-110 Mbp | 387,369 | - |

| | | | | |
|---|---|---|---|---|
| Common | 1 | chr6:30-410 Mbp | 60,793 | Overlaps with MHC (Major Histocompatibility Complex) region |
| Common | 2 | chr8:0-110 Mbp | 54,037 | Contains parts of the β-defensin gene clusters |
| Common | 3 | chr16:80-910 Mbp | 49,302 | Prone to structural variations and duplications |
| Common | 4 | chr8:10-210 Mbp | 48,197 | Associated with an immune response (BLK) |
| Common | 5 | chr16:0-110 Mbp | 44,329 | Prone to structural variations and duplications |
| Common | 6 | chr4:180-1910 Mbp | 43,360 | - |
| Common | 7 | chr7:0:110 Mbp | 42,340 | - |
| Common | 8 | chr3:0-110 Mbp | 41,827 | - |
| Common | 9 | chr9:0:110 Mbp | 41,775 | - |
| Common | 10 | chr4:0:110 Mbp | 41,037 | Associated with neurological development (HTT) |

**Table S2: Number of variants present in the HLA region for each population:** These variants were used to create the population-specific HLA genome graphs.

| Population | Number of Samples | Number of variants |
|---|---|---|
| AFR | 662 | 91,683 |
| AMR | 347 | 78,676 |
| EAS | 504 | 77,743 |
| EUR | 503 | 77,132 |
| SAS | 489 | 76,253 |

**Table S3: List of hypervariable nodes present in the population-specific HLA genome graphs:** The genomic positions correspond to the coordinates in hg38. Yes denotes the presence of a hypervariable node for the particular genome graph at that genomic position, and No denotes the absence of the same.

| Hypervariable Genomic Position | AFR HLA Genome Graph | AMR HLA Genome Graph | EAS HLA Genome Graph | EUR HLA Genome Graph | SAS HLA Genome Graph |
|---|---|---|---|---|---|
| 29,699,787 | Yes | No | No | Yes | No |
| 29,767,771 | Yes | Yes | No | Yes | Yes |
| 29,951,575 | Yes | Yes | No | Yes | No |
| 29,991,587 | No | Yes | Yes | Yes | No |
| 30,182,970 | Yes | Yes | No | No | No |
| 30,605860 | Yes | No | Yes | No | Yes |
| 31,340,514 | Yes | Yes | Yes | Yes | Yes |
| 31,340,587 | No | No | Yes | No | No |
| 31,412,380 | Yes | Yes | Yes | Yes | Yes |
| 31,412,383 | Yes | Yes | Yes | Yes | Yes |
| 32,211,594 | No | No | No | No | Yes |
| 32,223,881 | Yes | Yes | Yes | Yes | Yes |
| 32,303,314 | No | Yes | No | Yes | No |
| 32,332,596 | No | Yes | No | No | No |
| 32,492,191 | No | Yes | No | Yes | No |
| 32,589,513 | Yes | Yes | Yes | Yes | Yes |
| 32,964,843 | Yes | Yes | No | No | No |
| 33,127,190 | Yes | Yes | No | No | No |
| 33,127,191 | Yes | Yes | No | No | No |

### SUPPLEMENTARY FIGURES

**Figure S1:** Path-level representation of the most complex nodes in the 1KGP Genome Graph. Markers are placed on the edge that trace the reference path. The 12-degree nodes are present in chromosome 1, and the node IDs are *19298453 and 19298461*.

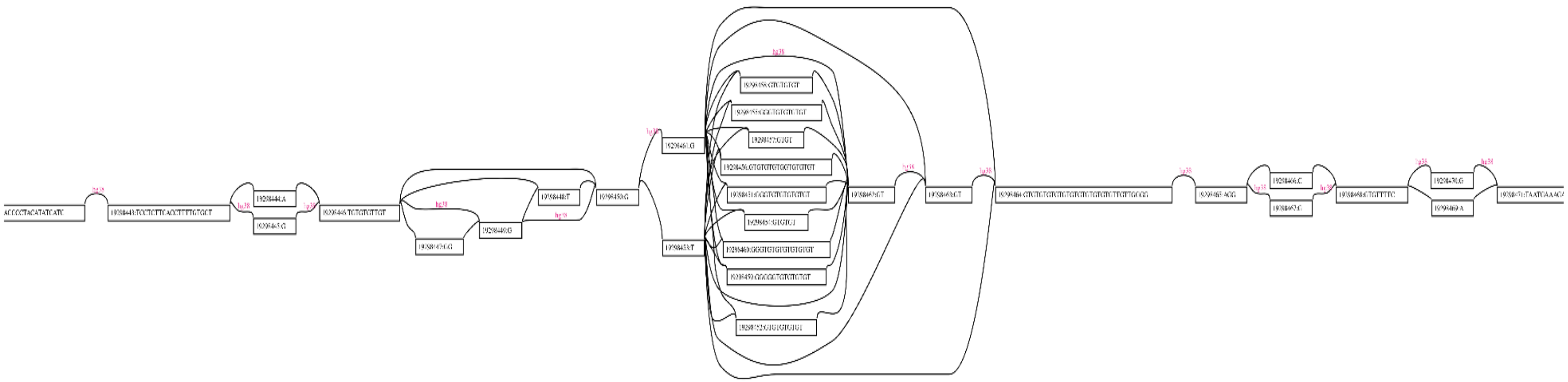

**Figure S2:** The median rate of recombination observed in each human chromosome.

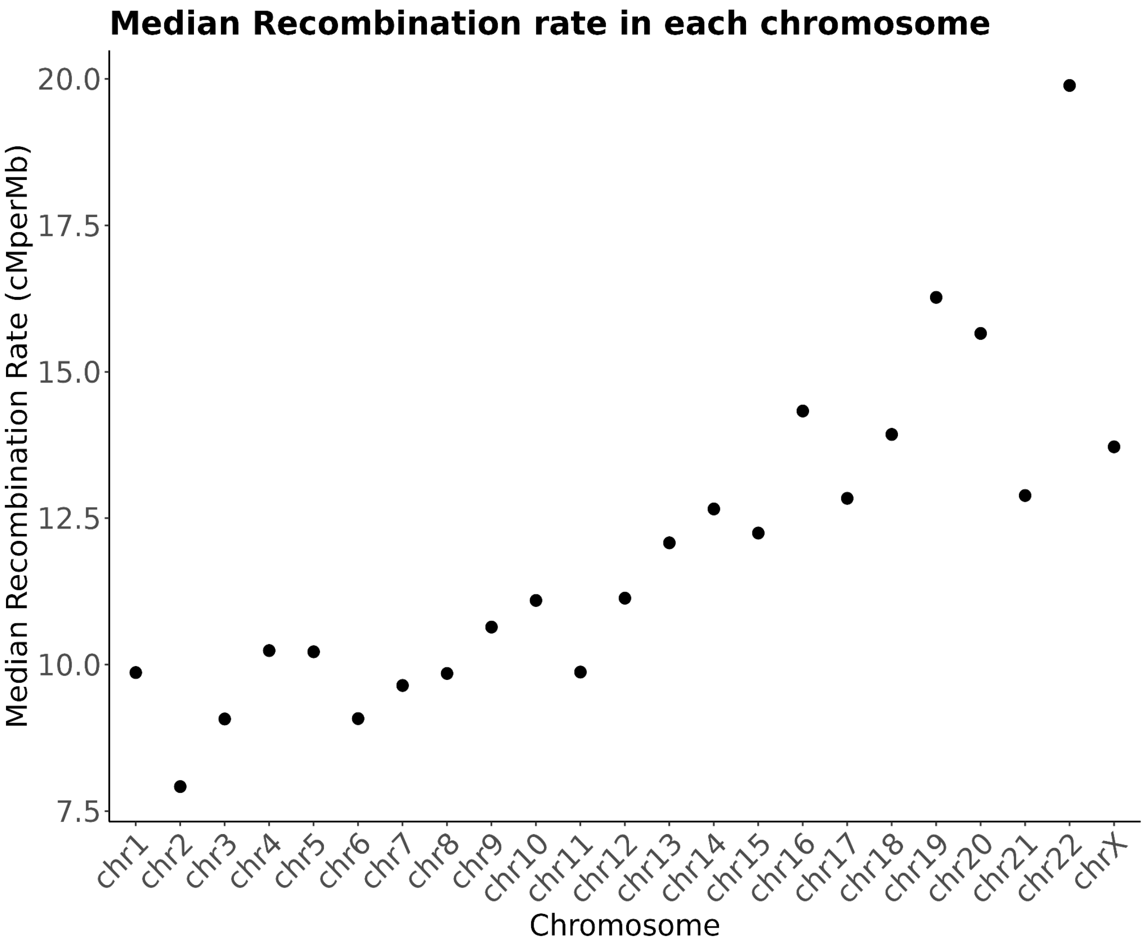
